# Predicting the specific substrate for transmembrane transport proteins using BERT language model

**DOI:** 10.1101/2022.07.23.501263

**Authors:** Sima Ataei, Gregory Butler

## Abstract

Transmembrane transport proteins play a vital role in cells’ metabolism by the selective passage of substrates through the cell membrane. Metabolic network reconstruction requires transport reactions that describe the specific substrate transported as well as the metabolic reactions of enzyme catalysis. In this paper, we apply BERT (Bidirectional Encoder Representations from Transformers) language model for protein sequences to predict one of 12 specific substrates. Our UniProt-ICAT-100 dataset is automatically constructed from UniProt using the ChEBI and GO ontologies to identify 4,112 proteins transporting 12 inorganic anion or cation substrates. We classified this dataset using three different models including Logistic Regression with an MCC of 0.81 and accuracy of 97.5%; Feed-forward Neural Networks classifier with an MCC of 0.88 and accuracy of 98.5%. Our third model utilizes a Fine-tuned BERT language model to predict the specific substrate with an MCC of 0.95 and accuracy of 99.3% on an independent test set.

## I. Introduction

Transmembrane transporter proteins play a vital role in cells’ metabolism by the selective passage of substrates across the cell membrane. This process includes the transportation of various compounds ranging from ions to macromolecules. The number of characterized transporters is small due to the difficult process of experimental characterization and determination of 3D structures. The large scale of sequencing today means that computational tools are necessary to facilitate the analysis of proteins in general, and membrane proteins, such as transporters, in particular.

The genome-scale reconstruction of a metabolic network (GENRE) is a key step in systems biology and synthetic biology. The process [1] builds a map to represent the geneprotein-reaction (GPR) association between the genes, the proteins which are the gene products, to the reaction that is carried out by the protein. This ideally covers metabolism, transport, regulation, and signaling. Typically a GENRE models metabolism quite well, and can assign GPR associations based on Enzyme Commission (EC) classification or Gene Ontology molecular function terms of the genes. The transport of substrates across membranes is modeled by transport reactions. For higher organisms, the models include the cellular compartments: the extracellular space, the cytosol, the mitochondrion, and sometimes the peroxisome. The prediction of GPR association for transport proteins, in particular, the specific substrate, or substrates, is not well-developed.

For homologs of known transporters, the standard applications of similarity search and HMM models of orthologous protein families have proven effective for annotation of transporters, as illustrated by TransportDB [2] and TCDB tools [3], TransATH [4], merlin [5]–[7], and Pantograph [8]. TranSyT [7] is a component of merlin 4.0 that utilises the mappings in TCDB and KEGG to determine substrates for transport reactions during the reconstruction of pathway networks by assigning TC identifiers, GO terms, and ChEBI terms to sequences.

For *de novo* prediction, where there are at best only remote homologs known, there is the work at the labs of Gromiha [9]– [11], Helms [12]–[14], Zhao [2], [15], and Butler [4], [16]–[18] and sporadic contributions from teams working in machine learning [19]–[22]. The best tools for classifiying substrate classes, based on the performance measures in their papers, are TrSSP [15] and FastTrans method [22], both on seven substrate classes, and *TooT-SC* [17], [18] on eleven substrate classes.

There is no previous work to our knowledge on predicting specific substrates *de novo* other than work predicting ion channels for sodium, potassium, and calcium ions [23] and work mining the scientific literature for a GENRE of human [24] that includes transport reactions.

Given the advances in representations of protein sequences, such as BERT language models, techniques from deep learning, and the growing number of sequences available in UniProt, we revisit the problem of predicting the specific substrate of a transmembrane transport protein.

The eleven classes for the *TooT-SC* dataset were constructed [25] using the substrate class descriptions in ChEBI and merging subclasses to obtain classes with sufficient number of sequences in Swiss-Prot to act as a dataset for machine learning. From [25, Table I and Table VI] we can see several classes with sufficient number of sequences to warrant consideration as starting points for specific substrates, namely

**TABLE I:** Four extracted datasets

| Dataset | Size | Trainset | Testset | # Classes |
| --- | --- | --- | --- | --- |
| UniProt-ICAT-100 | 4,112 | 3,289 | 823 | 12 |
| UniProt-ICAT-60 | 1,429 | 1,143 | 286 | 11 |
| SwissProt-ICAT-100 | 2,140 | 1,712 | 428 | 11 |
| SwissProt-ICAT-60 | 1,098 | 878 | 220 | 11 |

The number of sequences and classes in each of the four datasets. Each dataset has been divided into a training set and test set with 80%-20% ratio randomly and stratified.

1. CHEBI:15377 Water with 26 sequences
2. CHEBI:36915 Inorganic cation with 603 sequences
3. CHEBI:24834 Inorganic anion with 103 sequences
4. CHEBI:25696 Organic anion with 107 sequences

On these classes *TooT-SC* achieved an MCC of 1.00, 0.83, 0.83, and 0.81 respectively during independent testing.

In this paper, we introduce an automated method to construct datasets of transporter proteins and their exact corresponding substrates using Gene Ontology and `ChEBI` annotations. We constructed four datasets of inorganic cation and anion transporters from UniProt and Swiss-Prot using this method. We utilized a BERT language model for protein sequences to classify the proteins based on their specific substrate. We implement three classification models of Logistic Regression (LR), Feed-forward Neural Network and Fine-tuned separately. Our best classifier predicts the specific substrate of proteins with an MCC of 0.95 and accuracy of 99.3% during independent testing for our UniProt-ICAT-100 dataset with 12 substrate classes and 4,112 sequences.

## II. BERT Language Model for Proteins

Proteins are represented as a concatenation of amino acids. As such they share parallels in function and structure with natural languages [26]. Hence it would appear feasible to apply natural language processing (NLP) techniques to protein sequences. Transformer Neural Networks (Transformers) [27] have had a profound effect in NLP. Autoencoders, such as BERT (Bidirectional Encoder Representations from Transformers) [28] are stackable models trained by corrupting the input tokens and attempting to reconstruct the original phrase — that is, using a Masked Language Model (MLM) and Next Sentence Prediction (NSP) for pre-training bidirectional transformers [27]. They are most frequently used to create vector representations for subsequent tasks such as classification. The BERT language model generates representations depending on the sequence, and each amino acid has a unique meaning in each protein. This contrasts with context-free word2vec approaches such as prot2vec [29], which build a static representation of each amino acid in a protein.

Recent work on ProtTrans [30] studied six Transformer-based architectures to predict the secondary structure and subcellular localization. One model that performed well is ProtBERT-BFD [30], a pre-trained BERT model on a huge corpus of protein sequences from the BFD database (https://bfd.mmseqs.com) that employs the MLM task solely. The ProtBERT-BFD language model is composed of thirty layers of sixteen attention head transformer encoders with a total of 420 million parameters. Each amino acid is represented as a 1024-dimension vector in this language model.

BERT consists of two phases: pre-training and fine-tuning. The BERT model is trained without supervision on massive volumes of unlabeled data during pre-training using MLM and NSP tasks [28]. On the other hand, fine-tuning is the process of initializing the model with pre-trained parameters and updating its parameters using labeled data from subsequent tasks via an additional classifier [28]. The BERT model can be used to extract frozen or Fine-tuned representations. The frozen representation comprises features from a pre-trained BERT model that have not been updated, whereas the Fine-tuned representation has been Fine-tuned to the task at hand on a smaller labeled dataset [28]

Here we use the frozen pre-trained ProtBERT-BFD model to encode the protein sequences as well as fine-tuning the model for the classification task.

## III. Dataset Construction

A lack of research in predicting specific substrates in transporter activities results in absence of a dataset of transmembrane transport protein sequences with their specific substrate. We develop an automatic method to construct a dataset of transmembrane transport proteins and their specific substrate. The sequences come from the UniProt KB [31] and the Swiss-Prot subset. The substrates are identified using the ChEBI (Chemical Entities of Biological Interest) ontology [32] and the connection [33] is made to the protein sequence using the GO (Gene Ontology) [34], [35] annotations in UniProt.

We apply the algorithm to the above `ChEBI` terms

1. CHEBI:36915, Inorganic cation
2. CHEBI:24834, Inorganic anion

and the GO terms

1. GO:0022890, Inorganic cation transmembrane transporter activity (MF)
2. GO:0015103, Inorganic anion transmembrane transporter activity (MF)
3. GO:0003824, Catalytic activity (MF)

to differentiate between transporters and enzymes.

We create four datasets:

1. UniProt-ICAT-100, the algorithm applied to UniProt with CD-HIT at 100 percent identity for sequences;
2. Swiss-Prot-ICAT-100, the algorithm applied to Swiss-Prot with CD-HIT at 100 percent identity for sequences;
3. UniProt-ICAT-60, the algorithm applied to UniProt with CD-HIT at 60 percent identity for sequences; and
4. Swiss-Prot-ICAT-60, the algorithm applied to Swiss-Prot with CD-HIT at 60 percent identity for sequences.

To construct a dataset of transmembrane transport protein sequences with their specific substrate, a mapping, called C2GO, between `ChEBI` ontology and GO annotation is required. The three main steps are:

1. Build C2GO map from `ChEBI` leaf terms;
2. Build C2GO map to GO leaf terms; and
3. Find sequences with related GO terms and label with the corresponding `ChEBI` term.

The following sections describe each step.

### A. C2GO map from ChEBI leaves

The first step to label transporter protein sequences is to find the carried substrates from `ChEBI` ontology. As mentioned in the previous section, the dataset is built based on inorganic cation and inorganic anion substrates. To find these substrates, first, all `ChEBI` terms in the Directed Acyclic Graph (DAG) of ‘CHEBI:36915’ (inorganic cation), C_cation, and ‘CHEBI:24834’ (inorganic anion), C_anion, have been retrieved from ‘CHEBI.obo’ file. To attain the specific substrates, the leaf `ChEBI` terms in the C_anion and C_cation have been extracted according to `ChEBI` ontology, (C_leaf).

Next, the corresponding GO terms to each `ChEBI` term in C_leaf are retrieved from the ‘GO-Plus.OWL’ file. Each `ChEBI` term and its corresponding GO term is collected to build initial `ChEBI` to GO mapping, (C2GO). To capture all the GO terms corresponding to a specific `ChEBI` term, we find the DAG of each GO term in Gene Ontology and add it to the same `ChEBI` key as the father in the C2GO map. This process is described in Algorithm 1.

#### Algorithm 1

C2GO map from `ChEBI` leaf terms

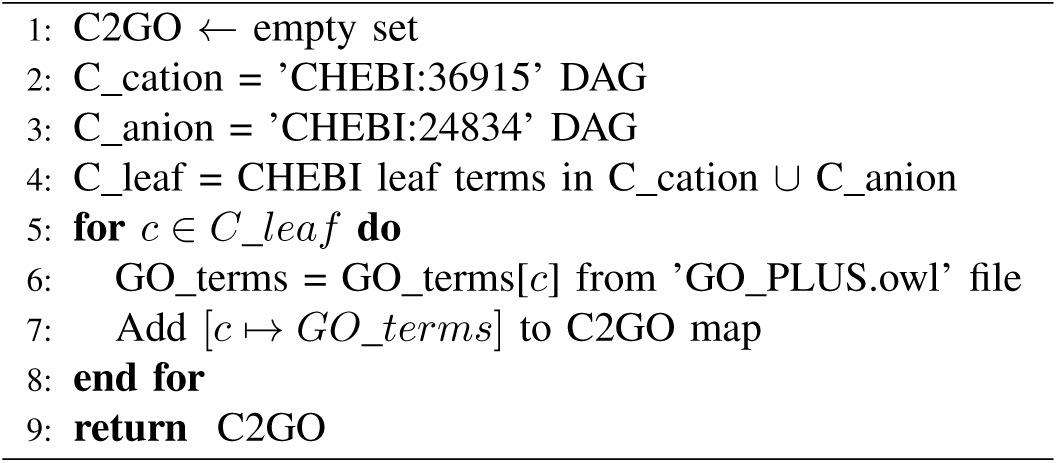

### B. C2GO map from GO leaves

We utilize the Molecular Function aspect of the Gene Ontology graph with the `is_a` relationship between the GO terms from the ‘GO-basic.OBO’ file. First, the GO terms in the DAG of ‘GO:0022890’ (inorganic cation transmembrane transporter activity) and ‘GO:0015103’ (inorganic anion transmembrane transporter activity) are retrieved from the graph, (GO_ICAT).

Next, the GO leaf terms according to Gene Ontology are extracted from this set to build GO)_ICAT leaf.

The GO terms ‘GO:0022890’ inorganic cation transmembrane transporter activity, ‘GO:0015103’ inorganic anion transmembrane transporter activity and ‘GO:0003824’ catalytic activity (MF) have common subtrees in the ontology. To differentiate between transporters and enzymes, the enzyme sequences are excluded from the dataset. To remove catalytic activity sequences from GO_ICAT_leaf set, DAG for ‘GO:0003824’ (catalytic activity), GO_catalytic is extracted from Gene Ontology. After GO leaf terms for GO_catalytic have been removed from GO_ICAT leaf inorganic cation and anion set, each GO_term in GO_ICAT leaf corresponding `ChEBI` term has been retrieved from the ‘GO-Plus.OWL’ file.

To exclude `ChEBI` terms unrelated inorganic cation and anion transporters, the extracted `ChEBI` terms from this step have been examined with the C_cation or C_anion sets. As the last step refined `ChEBI` terms and their corresponding GO terms have been added to the C2GO map. This process has been presented in Algorithm 2.

#### Algorithm 2

C2GO map from GO leaf terms

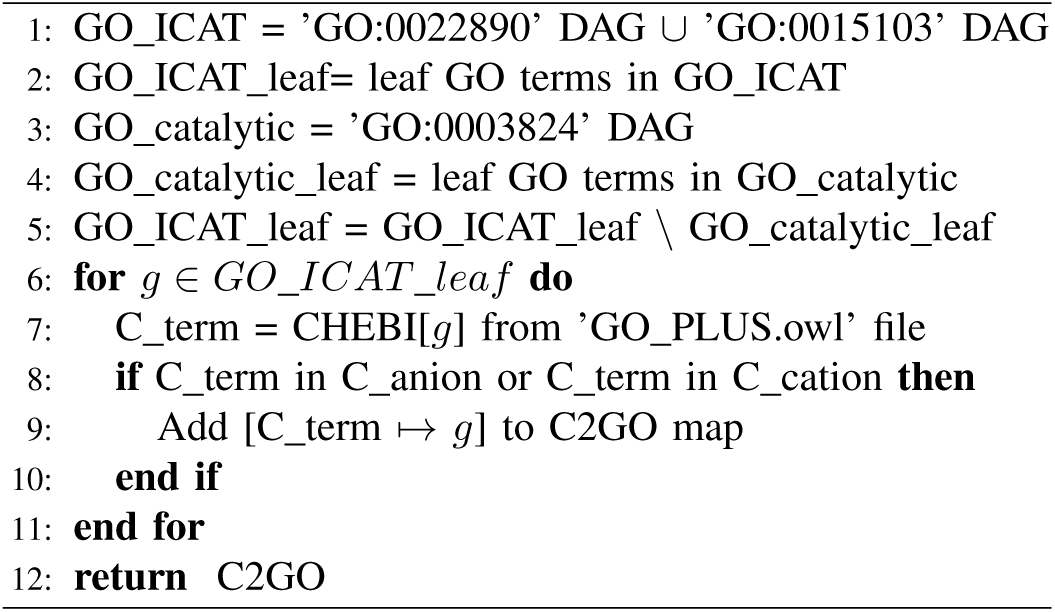

### C. Map Refinement and Labeling Sequences

In the Gene Ontology, there are GO terms with more than one `ChEBI` term associated with them. To improve the accuracy of the map, we decided to remove non-leaf `ChEBI` terms from GO terms with multiple `ChEBI` mappings.

Next, we delete `ChEBI` terms that have children in the mapping. Therefore, for all `ChEBI` terms in the C2GO keys, if children of the `ChEBI` term are present in the C2GO map, the `ChEBI` term itself is removed from the map. This process is described in Algorithm 3.

#### Algorithm 3

Refining C2GO map

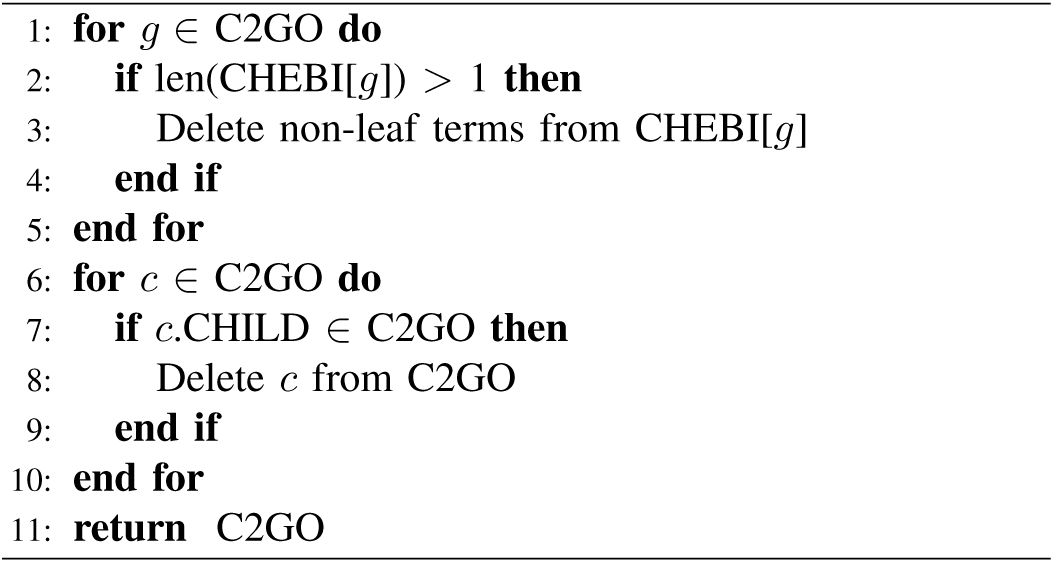

The final step is to label protein sequences according to the produced C2GO map. In this step we download the proteins with transmembrane transporter activity and the evidence at the protein level from UniProt with the following query:

⟨goa:(“transmembrane transporter activity [22857]”) existence:”Evidence at protein level [1]”⟩ and Swiss-Prot with:

⟨goa:(“transmembrane transporter activity [22857]”) existence:”Evidence at protein level [1]” reviewed:yes⟩

For each sequence in UniProt (or Swiss-Prot) Dataset, label the sequences with the `ChEBI` term if the sequences have common GO terms with the corresponding `ChEBI` term from the C2GO map. Finally, we remove sequences with more than one label or sequences without any label. This process is described in Algorithm 4.

#### Algorithm 4

Labeling Data

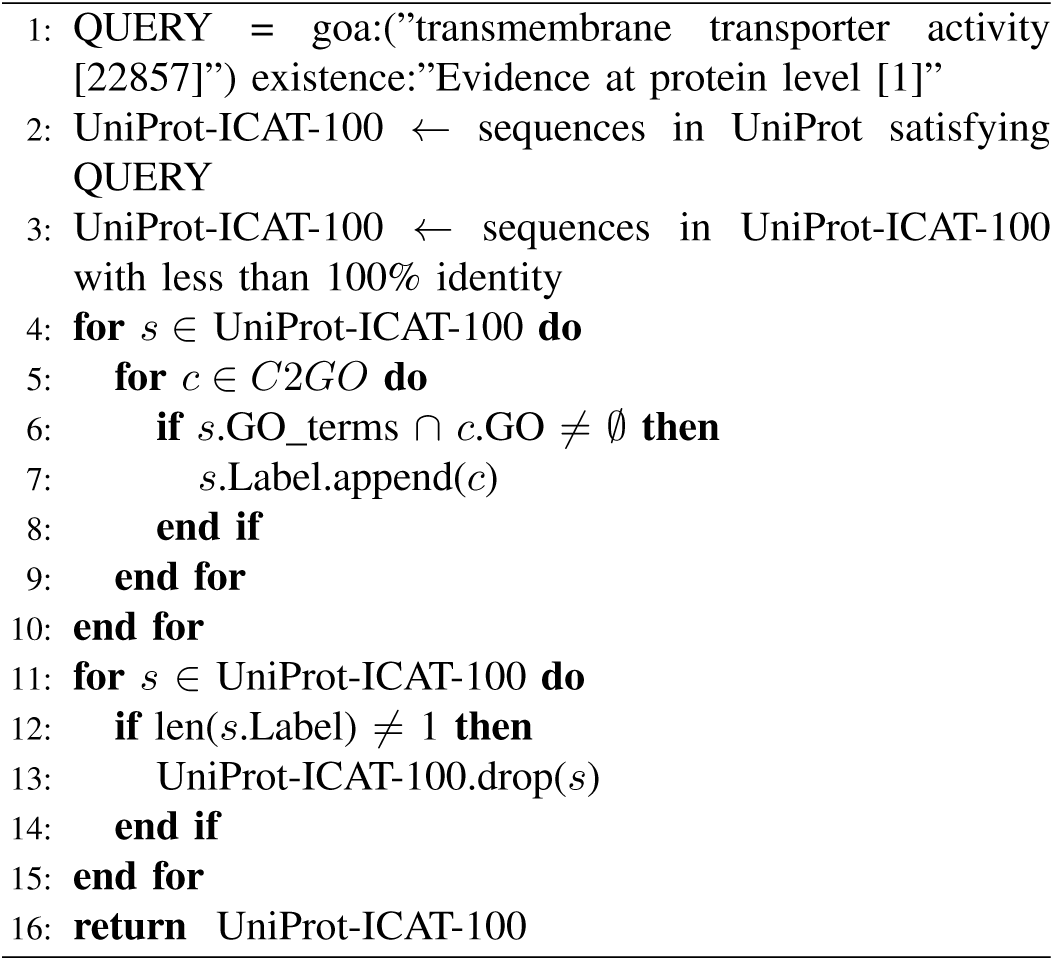

### D. Identity Reduction

We removed sequences with more than 100% and 60% pairwise sequence identity separately using the CD-HIT program [36]. A threshold of 10 sequences in each class has been applied and substrate classes with less than 10 sequences are eliminated from the datasets. The final datasets and the substrate classes are presented in detail in Table II.

**TABLE II:** Size of substrate classes for each dataset

| Class | CHEBI | Substrate | UP-ICAT-100 | UP-ICAT-60 | SP-ICAT-10 | SP-ICAT-60 |
| --- | --- | --- | --- | --- | --- | --- |
| 0 | CHEBI:24636 | proton | 1307 | 600 | 883 | 484 |
| 1 | CHEBI:29108 | calcium(2+) | 868 | 210 | 350 | 154 |
| 2 | CHEBI:29103 | potassium(1+) | 726 | 193 | 299 | 146 |
| 3 | CHEBI:17996 | chloride | 458 | 129 | 177 | 75 |
| 4 | CHEBI:29101 | sodium(1+) | 429 | 123 | 235 | 99 |
| 5 | CHEBI:16189 | sulfate | 98 | 49 | 38 | 27 |
| 6 | CHEBI:29105 | zinc(2+) | 80 | 51 | 67 | 47 |
| 7 | CHEBI:28938 | ammonium | 55 | 28 | 36 | 23 |
| 8 | CHEBI:17632 | nitrate | 28 | 18 | 27 | 18 |
| 9 | CHEBI:29033 | iron(2+) | 24 | 16 | 16 | 13 |
| 10 | CHEBI:35780 | phosphate ion | 22 | 12 | 12 | 12 |
| 11 | CHEBI:49552 | copper(1+) | 17 | 9 | 7 | 6 |
| Total |  |  | 4,112 | 1,429 | 2,140 | 1,098 |

The 12 specific substrates and the number of sequences for that substrate in each of the four datasets: UniProt-ICAT-100, Swiss-Prot-ICAT-100, UniProt-ICAT-60, and Swiss-Prot-ICAT-60 respectively. Numbers in *italic* are below the threshold of 10 sequences. Their substrate class is not included in experiments with that dataset.

## IV. Methods

We implement three different approaches based on the ProtBERT language model to classify protein sequences. As the first method, we use a frozen ProtBERT-BFD language model to extract protein representations followed by a Logistic Regression algorithm as a classifier. For the second method, similar to the first one, we use frozen ProtBERT-BFD representations followed by a Feed-forward Neural Network to classify sequences. The third approach is to Fine-tune the same ProtBERT-BFD model with a linear layer and a softmax function for classification.

As Table I presents, each dataset has been divided into the training set and independent test set with a ratio of 80% to 20%. For Logistic Regression classification, a 5-fold cross-validation has been performed on the training set. Then the dataset is trained on the full training set and tested on the independent test set. The other two methods are validated after each epoch of training process

### A. BERT Representation

To produce representations of protein sequences we used the ProtBERT-BFD pre-trained model publicly available on the HuggingFace website [37]. In this approach, we freeze the ProtBERT-BFD architecture and extract encodings of proteins from the last hidden layer. For each sequence, we collect the last hidden layer representation of the first token [CLS] which contains the encodings of all the amino acids in the sequence. The maximum length of sequences has been set to 1024 as the pre-trained model and larger sequence lengths are truncated. The ProtBERT-BFD representations are passed to a Logistic Regression and a Feed-forward Neural Network separately for the classification task.

### B. Logistic Regression

We chose Logistic Regression (LR) as a basic classification technique [38]. A multi-class classification has been implemented for 5-fold cross-validation and independent testing. This model is implemented using scikit-learn python package [39].

### C. Feed Forward Neural Network

As the second classification method, we trained a Feed-forward Neural Network with 100 units in one hidden layer for 100 epochs [40]. The input layer size is set to 1024, as the length of ProtBERT-BFD produced representations, and the output layer size is set to the number of classes (11 or 12 depending on the dataset) available in the corresponding dataset. The network utilizes the Cross entropy loss function and Stochastic Gradient Descent (SGD) as an optimizer [41]. The batch size is set to 1 sequence and the learning rate is 0.05 for UniProt-ICAT-60, SwissProt-ICAT-100, SwissProt-ICAT-60, and 0.005 for UniProt-ICAT-100 due to the dataset size.

### D. Fine-tuned BERT language model

In this method, pre-trained ProtBERT-BFD has been Fine-tuned on the extracted datasets separately to update the model weights for the specified classification task. Our model includes a single-layer Feed-forward network followed by a softmax function to predict the probabilities of each class in the dataset. The configuration for the network is as follows; Cross entropy loss function and Adam optimizer are utilized with learning rate = 5 ×10^−5^ for batch size 1 [42]. The training process achieves acceptable results in 7 epochs. The implementations are applied using the Pytorch package and the pre-trained ProtBERT-BFD model is collected from the HuggingFace website [37], [43].

### E. Evaluation Metrics

The performance of each method on the training set was determined using five-fold cross-validation and reported the performance variations between the runs by computing the standard deviation. Five performance metrics were considered:

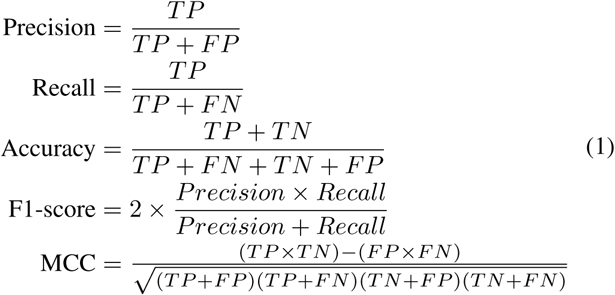

Where *TP* is the number of true positives, *TN* is the number of true negatives, *FP* is the number of false positives, and *FN* is the number of false negatives.

The Matthews Correlation Coefficient (MCC) is less influenced by imbalanced data and is arguably the best single assessment metric in this case [44]–[46]. The overall performance across all classes is the micro-average of the individual results [47] and we used the multi-class version of MCC [48].

## V. Results and Discussion

Five-fold cross-validation is performed for the using frozen BERT and Logistic Regression classifier and the Table III presents the overall results. For five-fold cross-validation on Logistic Regression classification method, the largest dataset UniProt-ICAT-100 achieves the highest MCC of 0.81. As the Table III presents for each dataset, the variation in performance is small, as shown by the standard deviation. The Logistic Regression model for independent testing is trained on the entire training set. It has outperformed the Logistic Regression models during cross-validation which are trained on a subset of the training set and tested by the validation set in all the metrics.

**TABLE III:** 5-fold cross-validation results for each dataset using frozen BERT and logistic regression classifier

| Dataset | Classes | Accuracy | Precision | Recall | F1-score | MCC |
| --- | --- | --- | --- | --- | --- | --- |
| UniProt-ICAT-100 | 12 | **0.848 ± 0.01** | **0.679 ± 0.05** | **0.561 ± 0.02** | **0.598 ± 0.02** | **0.808 ± 0.02** |
| UniProt-ICAT-60 | 11 | 0.727 ± 0.03 | 0.586 ± 0.08 | 0.430 ± 0.04 | 0.467 ± 0.05 | 0.631 ± 0.04 |
| SwissProt-ICAT-100 | 11 | 0.825 ± 0.01 | 0.628 ± 0.04 | 0.512 ± 0.04 | 0.542 ± 0.04 | 0.766 ± 0.02 |
| SwissProt-ICAT-60 | 11 | 0.723 ± 0.03 | 0.476 ± 0.06 | 0.395 ± 0.04 | 0.415 ± 0.05 | 0.617 ± 0.05 |

Figure 2 shows the validation performance in the training process for both Feed-forward Neural Network and Fine-tuned Prot-BERT using UniProt-ICAT-100. As 2a shows the Feed-forward Neural Network reaches maximum performance on validation set in epoch 89 with MCC of 0.900. Similarly, Figure 2b shows the learning curve maximum on epoch number 7 with MCC of 0.950.

**Fig. 1:**
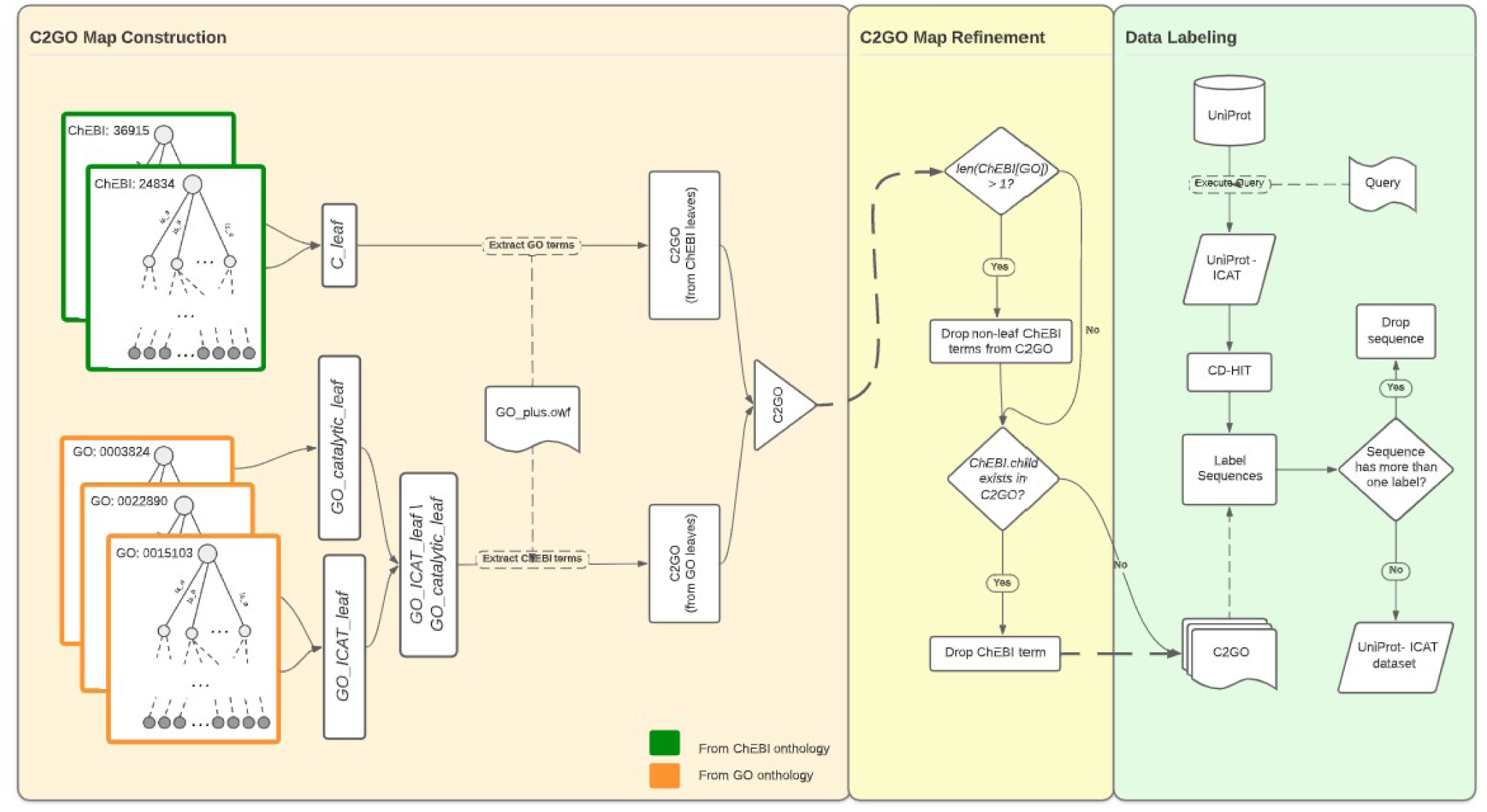
The process of constructing UniProt-ICAT-100 dataset: C2GO map Construction from Directed Acyclic Graph from Gene Ontology and CHEBI annotation, based on algorithms 1 and 2 (left). C2GO map is refined in two steps as algorithm 3 describes (middle). The data labeling is performed based on the extracted C2GO map based on algorithm 4 (right)

**Fig. 2:**
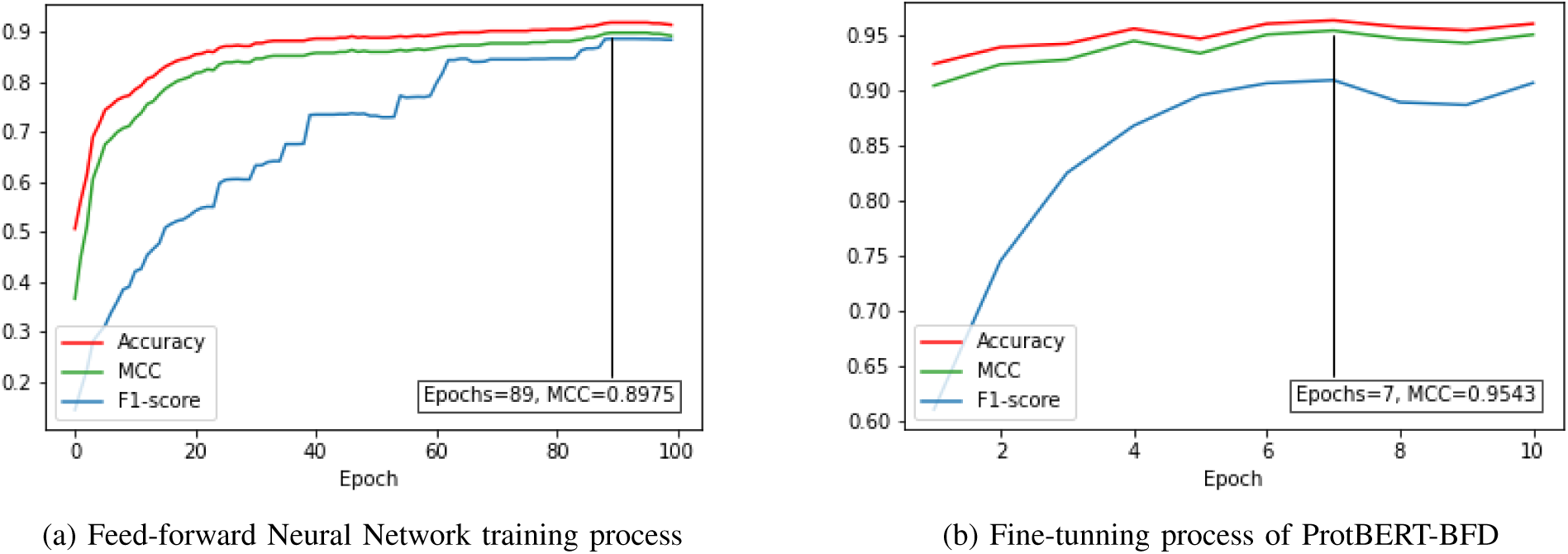
Camparison of validation performance in training process for two classification methods using UniProt-ICAT-100

Table IV presents the overall results of the independent testset for three methods implemented on each of the datasets. This table suggests the Fine-tuned model outperforms the other methods in all the metrics for four datasets. Table V presents detailed results for the fined-tuned ProtBERT-BFD model on UniProt-ICAT-100 dataset for each class in independent testset. Despite the small size in the dataset, Copper ion (`copper(1+)`) is predicted completely correct. Three classes of `zinc(2+)`, `iron(2+)` and `nitrate` have scored 100% in precision, and most of the classes achieve very strong MCC scores. The Figure 3 presents the confusion matrix for this results.

**TABLE IV:** Independent testset results comparison for Logistic Regression(LR), Feed-forward Neural Networks (FNN), and Fine-tuned BERT (FTB)

| Dataset | # Classes | Accuracy |  |  | Precision |  |  | Recall |  |  | F1-score |  |  | MCC |  |  |
| --- | --- | --- | --- | --- | --- | --- | --- | --- | --- | --- | --- | --- | --- | --- | --- | --- |
|  |  | LR | FNN | FTB | LR | FNN | FTB | LR | FNN | FTB | LR | FNN | FTB | LR | FNN | FTB |
| UniProt-ICAT-100 | 12 | 0.975 | 0.985 | **0.993** | 0.852 | 0.911 | **0.959** | 0.630 | 0.832 | **0.881** | 0.680 | 0.858 | **0.913** | 0.813 | 0.889 | **0.948** |
| UniProt-ICAT-60 | 11 | 0.952 | 0.963 | 0.982 | 0.734 | 0.794 | 0.899 | 0.415 | 0.597 | 0.868 | 0.442 | 0.664 | 0.903 | 0.640 | 0.726 | 0.867 |
| SwissProt-ICAT-100 | 11 | 0.971 | 0.981 | 0.991 | 0.839 | 0.895 | 0.951 | 0.509 | 0.748 | 0.879 | 0.548 | 0.783 | 0.907 | 0.785 | 0.861 | 0.936 |
| SwissProt-ICAT-60 | 11 | 0.950 | 0.956 | 0.979 | 0.723 | 0.759 | 0.886 | 0.425 | 0.595 | 0.757 | 0.465 | 0.598 | 0.802 | 0.616 | 0.682 | 0.850 |

**TABLE V:** Detailed results for classification of UniProt-ICAT-100 using Fine-tuned ProtBERT

| Substrate | Trainset | Validation | Testset | TP | FP | FN | TN | Accuracy | Precision | Recall | F1-Score | MCC |
| --- | --- | --- | --- | --- | --- | --- | --- | --- | --- | --- | --- | --- |
| proton | 837 | 209 | 261 | 245 | 17 | 16 | 545 | 0.960 | 0.935 | 0.939 | 0.937 | 0.908 |
| calcium(2+) | 555 | 139 | 174 | 157 | 12 | 17 | 637 | 0.965 | 0.929 | 0.902 | 0.916 | 0.893 |
| potassium(1+) | 465 | 116 | 145 | 136 | 15 | 9 | 663 | 0.971 | 0.901 | 0.938 | 0.919 | 0.901 |
| chloride | 293 | 73 | 92 | 86 | 9 | 6 | 722 | 0.982 | 0.905 | 0.935 | 0.920 | 0.910 |
| sodium(1+) | 274 | 69 | 86 | 75 | 14 | 11 | 723 | 0.970 | 0.843 | 0.872 | 0.857 | 0.840 |
| sulfate | 62 | 16 | 20 | 14 | 4 | 6 | 799 | 0.988 | 0.778 | 0.700 | 0.737 | 0.732 |
| zinc(2+) | 51 | 13 | 16 | 14 | 0 | 2 | 807 | 0.998 | **1.000** | 0.875 | 0.933 | 0.934 |
| ammonium | 35 | 9 | 11 | 11 | 1 | 0 | 811 | 0.999 | 0.917 | **1.000** | 0.957 | 0.957 |
| nitrate | 18 | 4 | 6 | 4 | 0 | 2 | 817 | 0.998 | **1.000** | 0.667 | 0.800 | 0.816 |
| iron(2+) | 15 | 4 | 5 | 2 | 0 | 3 | 818 | 0.996 | **1.000** | 0.400 | 0.571 | 0.631 |
| phosphate ion | 15 | 3 | 4 | 3 | 1 | 1 | 818 | 0.998 | 0.750 | 0.750 | 0.750 | 0.749 |
| copper(1+) | 11 | 3 | 3 | 3 | 0 | 0 | 820 | **1.000** | **1.000** | **1.000** | **1.000** | **1.000** |

Results for classification of each of the 12 specific substrates for a multi-class classifier trained on the training set of UniProt-ICAT-100 using Fine-tuned ProtBERT model and evaluated with the independent test set. Numbers in bold indicate perfect scores.

**Fig. 3:**
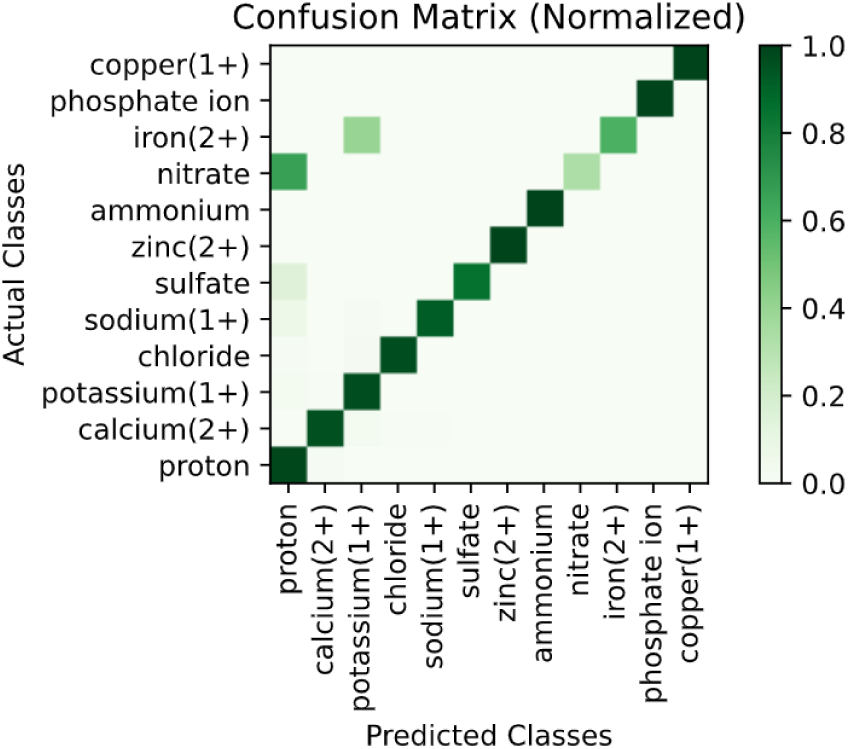
Normalized confusion matrix for classification results of the independent test using Fine-tuned BERT on UniProt-ICAT-100 dataset

From Table V we can see the number of classes has less effect on the dataset than the number of samples. This table shows that UniProt-ICAT-100 with 12 classes has better results in all the metrics than Swiss-Prot-ICAT-100 with 11 classes because of the higher number of sequences in this dataset.

## VI. Conclusion

This work introduces a BERT-based model for the classification of substrates carried by inorganic cation and anion transporters. By fine-tuning the ProtBERT model, we achieved a classifier to predict the specific inorganic ion with MCC of 0.95. The experiments show promising results compared to frozen BERT representation classified with Logistic Regression and Feed-forward Neural Networks. The use of CD-HIT to remove similar sequences is a suggested step in the methodology of bioinformatics machine learning, yet here it had a large impact on the number of sequences in many of the classes, and that impacted performance. Nevertheless, the results of this research suggest Prot-BERT language model’s potentials to investigate in other areas of protein classification research.

